# A Short Bout of High-intensity Exercise Alters Ipsilesional Motor Cortical Excitability Post-stroke

**DOI:** 10.1101/530857

**Authors:** Xin Li, Charalambos C. Charalambous, Darcy S. Reisman, Susanne M. Morton

## Abstract

**Background:** Acute exercise can increase motor cortical excitability and enhance motor learning in healthy individuals, an effect known as exercise priming. Whether it has the same effects in people with stroke is unclear.

**Objectives:** The objective of this study was to investigate whether a short, clinically-feasible high-intensity exercise protocol can increase motor cortical excitability in non-exercised muscles of chronic stroke survivors.

**Methods:** Thirteen participants with chronic, unilateral stroke participated in two sessions, at least one week apart, in a crossover design. In each session, they underwent either high-intensity lower extremity exercise or quiet rest. Motor cortical excitability of the extensor carpi radialis muscles was measured bilaterally with transcranial magnetic stimulation before and immediately after either exercise or rest. Motor cortical excitability changes (post-exercise or rest measures normalized to pre-test measures) were compared between exercise vs. rest conditions.

**Results:** All participants were able to reach the target high-intensity exercise level. Blood lactate levels increased significantly after exercise (*p* < 0.001, *d* = 2.85). Resting motor evoked potentials from the lesioned hemisphere increased after exercise compared to the rest condition (*p* = 0.046, *d* = 2.76), but this was not the case for the non-lesioned hemisphere (*p* = 0.406, *d* = 0.25).

**Conclusions:** High-intensity exercise can increase lesioned hemisphere motor cortical excitability in a non-exercised muscle post-stroke. Our short and clinically-feasible exercise protocol shows promise as a potential priming method in stroke rehabilitation.

## Introduction

Many stroke survivors have persistent motor function deficits despite receiving intensive rehabilitation^1,2^. These deficits lead to increased disability and decreased activity and participation^3,4^. Motor impairments post-stroke are associated with motor cortical excitability changes within the primary motor cortex (M1), which can be measured with transcranial magnetic stimulation (TMS). After stroke affecting the corticospinal pathway, motor evoked potentials (MEPs) from the lesioned hemisphere to the paretic upper extremity are decreased compared to the non-lesioned hemisphere^5,6^ and compared to healthy controls^6^. Furthermore, increases in lesioned hemisphere MEP amplitudes are associated with motor function improvement^5–7^. Therefore, interventions that increase motor cortical excitability within the lesioned hemisphere may be beneficial for motor recovery post-stroke.

Exercise can increase excitability broadly within the brain. That is, brain areas not directly involved in performance of the exercise show increased excitability following exercise^8^, an effect known as “exercise priming”. Exercise priming is potentially beneficial for stroke rehabilitation because excitability within brain regions controlling more impaired muscle groups or limbs might be modifiable by exercising intact limbs, thereby improving excitability within impaired brain areas without fatiguing impaired limbs or being limited by an inability to exercise very severely impaired limbs. Rehabilitation treatments then may capitalize on the increased cortical excitability, potentially producing enhanced motor recovery.

However, most work showing the effects of exercise on motor cortical excitability has been done in young, healthy adults. In this population, a single bout of lower extremity cycling can increase motor cortical excitability of a non-exercised upper extremity muscle^9–11^. Behaviorally, a single bout of high-intensity cycling has been shown to increase retention of learning an upper extremity motor task in healthy individuals, compared to controls that do not exercise^12,13^. Moreover, increases of motor cortical excitability after high-intensity cycling were positively correlated with the degree of retention of a motor sequence learning task^14^, suggesting a potential mechanism of behavioral gains from exercise through neurophysiological increases in motor cortical excitability.

In stroke, studies of the effects of a single bout of acute exercise are limited. A single bout of low-to-moderate intensity cycling failed to induce motor cortical excitability changes in a hand muscle in chronic stroke survivors^15^. Since high-intensity exercise is best at enhancing retention in healthy individuals^16^, the exercise intensity used in that study may not have been enough to induce changes in the stroke-affected brain. Utilizing high-intensity lower extremity exercise, Nepveu et al^17^ found that exercise increased retention of a time-on-target hand motor task. There was also a significant change in the relationship between intracortical inhibition in the lesioned and non-lesioned hemispheres compared to before exercise^18^. Another study reported latency changes in the lesioned hemisphere MEP after high-intensity interval training, but not after moderate-intensity exercise producing the same amount of work^19^.

The previous studies together suggest that high-intensity exercise, rather than low or moderate-intensity exercise, may induce changes of motor cortical excitability in the stroke-affected brain. However, the exercise protocols used in these studies are 20-25 minutes long. Although it is clear from these studies that stroke survivors can complete high-intensity exercise of this caliber, it is, however, probably unrealistic in a clinical setting, given that a typical physical therapy session is about 30-60 minutes. Incorporating this type of exercise priming protocol into a therapy session would significantly decrease the time allotted for rehabilitation treatment, and could potentially fatigue patients, thus limiting their ability to participate in more intensive therapy. In addition, longer exercise sessions may not be feasible for individuals with more than mild motor impairments post-stroke. Therefore, we aimed to determine whether a shorter bout of high-intensity exercise priming can enhance motor cortical excitability in individuals with stroke, including individuals with moderate and severe motor impairments. To that end, we used a 5 minute high-intensity exercise protocol that we have found to be effective in producing significant lactate level changes, a marker of exercise intensity, in people with chronic stroke^20^. We hypothesized that motor cortical excitability of non-exercised upper extremity muscles would increase bilaterally after a short bout of high-intensity lower extremity exercise.

## Methods

### Participants

Thirteen participants with chronic (> 6 months), unilateral stroke affecting the corticospinal tract participated in the study. They all went through a sensorimotor evaluation to determine their functional status, and brain MRI or CT scan reports were obtained to verify stroke type and lesion location. All participants were able to walk on a treadmill continuously for 5 minutes without assistance. Exclusion criteria included multiple strokes affecting both hemispheres, any cerebellar lesion, or any other significant neurological, cardiovascular or musculoskeletal condition besides stroke. Individuals with contraindications to TMS and those taking medications or substances affecting the central nervous system were also excluded. The experimental protocol was approved by the University of Delaware Institutional Review Board and conformed to the Declaration of Helsinki. All participants gave written informed consent. Participant demographic information is provided in Table 1.

**Table 1.**
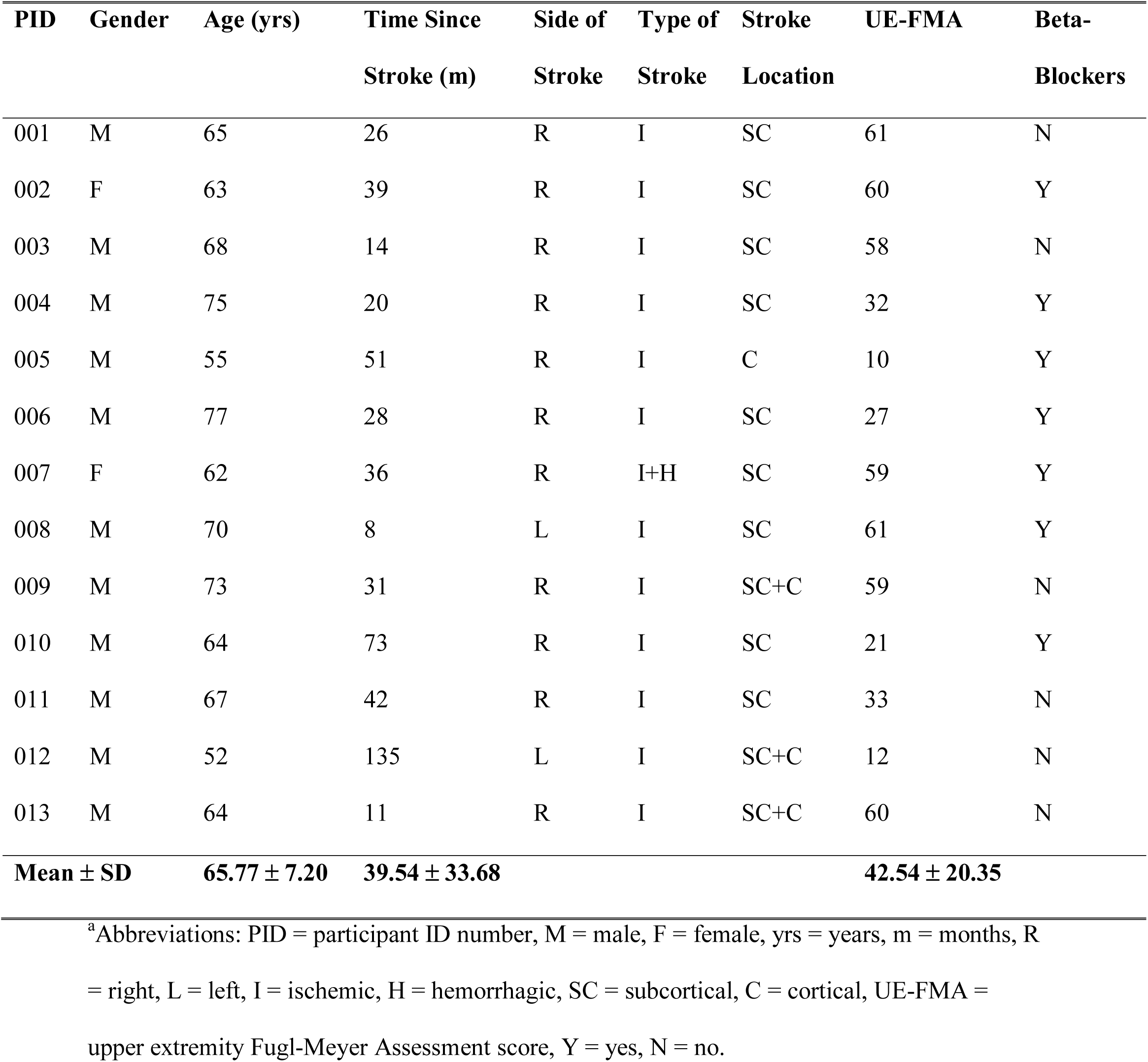
Participant demographic information^a^.

### General Paradigm

Participants completed two testing sessions separated by at least one week. They performed a short bout of high-intensity exercise priming (i.e. fast treadmill walking) in one session and sat quietly in the other session. Before and immediately after exercise or rest, motor cortical excitability of the extensor carpi radialis (ECR) muscles was measured bilaterally with TMS. Note that the ECR muscles were not directly involved in the high-intensity exercise (i.e., an upper extremity muscle), therefore, any changes of motor cortical excitability observed would be from the broad excitability changes induced by exercise, rather than a direct result of repeated activation of the muscle during exercise.

### Electromyography

Electromyography (EMG) was collected using a 10-channel system with double differential surface electrodes with an integrated ground (Motion Lab Systems, Inc., Baton Rouge, LA). EMG electrodes were placed over the ECR muscle bellies bilaterally, longitudinally following the direction of the muscle fibers. A ground electrode was placed over the lateral epicondyle of the humerus. EMG data were collected with a sampling rate of 5000 Hz and online low-pass filtered at 2000 Hz.

### Transcranial Magnetic Stimulation

For TMS, participants were seated and instructed to completely relax their arms with the elbows flexed over a pillow or armrests. A 70 mm diameter figure-of-eight coil was used in conjunction with Magstim 200^2^ and BiStim^2^ electromagnetic stimulation units (Magstim, Ltd., Wales, UK) for single-pulse and paired-pulse TMS measures, respectively. Signal 6.03 software (Cambridge Electronic Design, Ltd., Cambridge, UK) was used to control and trigger the magnetic stimulator through a 16-bit data acquisition unit (Micro 1401-3, Cambridge Electronic Design, Ltd., Cambridge, UK), and to collect and store EMG data for offline analysis.

The coil was held tangential to the scalp at a 45° angle to the mid-sagittal line to induce a posterior-to-anterior current in the upper extremity motor strip^21–23^. The “hot spot” for the ECR muscle was defined as the spot that produced the biggest and most consistent peak-to-peak MEP amplitudes at a given stimulus intensity, and marked carefully on the scalp for use throughout the session. Identical procedures were used to identify the “hot spot” in both sessions. In cases where no MEPs could be elicited from the lesioned hemisphere at 100% of the maximum stimulator output (MSO), we used a symmetrical “hot spot” to the non-lesioned hemisphere in reference to the vertex. Resting motor threshold (RMT) was defined as the lowest stimulus intensity that produced at least 5 out of 10 MEPs in the contralateral ECR muscle with a peak-to-peak amplitude over 50 μV^24^. Ten MEPs were collected from each hemisphere at 120% RMT before and after exercise or rest.

During the exercise session only, we also assessed short-interval intracortical inhibition (SICI). This measure was selected because SICI has consistently been shown to change in response to exercise priming in healthy individuals^10,25,26^. With the participant relaxed, a conditioning stimulus (CS) was delivered at an intensity of 90% RMT, followed by a test stimulus (TS) delivered at 130% RMT, with the inter-stimulus interval (ISI) set at 2.5 ms^27^. Ten trials each of the TS alone and of the paired CS-TS, presented in random order, were delivered to each hemisphere.

### Exercise / Rest Protocol

The rest protocol (rest session only) consisted of 10 minutes of quiet sitting. The exercise protocol (exercise priming session only) used was similar to that described in our previous work^20^. Briefly, participants completed 5 minutes of high-intensity walking on an instrumented treadmill (Bertec Corp., Columbus, OH). We defined high-intensity as exercise reaching 70% − 85% of the age-adjusted maximum heart rate (HR) (220 – age)^28^, or, for participants taking beta-blocker medications, reaching a rate of perceived exertion (RPE) of 13 - 15 on the Borg scale (scale of 6 - 20)^29,30^. HR was tracked during walking with a wireless HR monitor (Polar Electro Inc., Lake Success, NY) placed around the chest with data telemetered to a wristwatch. HR and RPE were recorded every 15 and 30 seconds, respectively. To reach and maintain the target HR or RPE throughout exercise, treadmill speed was adjusted around the participant’s fastest comfortable speed, which was determined at the beginning of data collection, i.e., before any TMS measurements. This exercise protocol has previously shown to be effective and tolerable for participants with stroke^20^. A harness and a front handrail were used during all walking procedures to ensure safety. Finger-tip blood lactate was measured using a portable Lactate Plus Meter (Nova Biomedical, Waltham, MA) before and immediately after exercise as an additional measure of exercise intensity^31^. EMG of the ECRs was collected throughout the exercise and analyzed offline to ensure that participants were not exercising the wrist muscles during walking.

### Data Analysis

EMG data were analyzed in Signal (Cambridge Electronic Design, Ltd., Cambridge, UK) and custom written software in MATLAB (MathWorks, Inc., Natick, MA). All raw EMG data were demeaned, and any gain removed. For all trials, EMG was notch-filtered at 60 Hz with a 2^nd^ order Butterworth filter to remove electrical noise, and trials with baseline (10 - 60 ms before stimulus artifact) peak-to-peak EMG exceeding 30 μV were discarded, as this could indicate the participant was not at rest. Peak-to-peak MEP amplitudes were calculated for each trial at 120% RMT and averaged over all trials. Resting MEP amplitudes were normalized as a ratio of that obtained during the post-test (for exercise or rest) to that obtained during the pre-test. This allowed us to compare changes of motor cortical excitability in individuals with widely varying degrees of impairment. SICI was calculated as the ratio of the mean peak-to-peak MEP amplitude of CS-TS trials to TS-only trials. Thus, a ratio value of 1 would indicate no intracortical inhibition, whereas values of < 1 would indicate inhibition.

Time at target intensity (either HR or RPE) was averaged across all participants^20^ and used as verification of adequate exercise intensity being reached. Blood lactate levels, compared before and after exercise, were used as a secondary marker for exercise intensity.

### Statistical Analysis

Statistical analyses were performed in IBM SPSS Statistics 24 (IBM Corp., Armonk, NY). Paired t-tests were applied to compare normalized resting MEPs (exercise vs rest session) and SICI ratios (pre to post exercise). However, for some measures for some individuals, we were unable to obtain MEPs from the lesioned hemisphere, thus reducing the sample size. Hence, when sample size was low or when data were not normally distributed, nonparametric statistics were used instead (Wilcoxon signed-rank test). Paired t-tests were also used to compare lactate levels before vs. after exercise.

## Results

All participants completed the experiment with no adverse events. All group data are reported as mean ± SEM unless otherwise stated.

### Exercise intensity

All participants were able to reach the target exercise level, and the average time spent at the target level was 66.15% ± 4.35% (range 35% - 85%) of the total exercise time. The secondary marker for exercise intensity, blood lactate levels, increased significantly post-exercise (4.28 ± 0.36 mmol/L) compared to pre-exercise (1.68 ± 0.11 mmol/L), *t*(12) = −7.53, *p* < 0.001, *d* = 2.85.

### Valid TMS data and baseline motor cortical excitability between sessions

We were able to obtain complete datasets from the non-lesioned hemisphere of all participants. From the lesioned hemisphere, during the exercise session, we were able to obtain RMTs from 8 participants (62%) and MEP amplitudes at 120% RMT and SICI from 6 participants (46%). During the rest session, we were able to obtain RMTs, MEP amplitudes and SICI from 6 participants (46%). Baseline TMS measures were similar across the two sessions (all *p* > 0.17).

### Resting MEP amplitude changes after exercise / rest

In Figure 1, we show traces of resting MEPs from one exemplar individual. Note that the MEP from the lesioned hemisphere had a larger peak-to-peak amplitude following exercise (compare thick black trace to thin gray trace, bottom panel, right), while it remained approximately the same size after rest (upper panel, right). This observation was confirmed in the statistical comparison of the entire group. Figure 2 shows group normalized resting MEP amplitudes after exercise and rest for the non-lesioned and lesioned hemispheres. Here it can be seen that excitability within the lesioned hemisphere increased more after exercise (1.66 ± 0.24) than rest (1.23 ± 0.30), *Z* = 1.99, *p* = 0.046, *d* = 2.76. There were no differences between exercise (1.33 ± 0.19) and rest (1.17 ± 0.16) conditions in the non-lesioned hemisphere, *t*(12) = 0.86, *p* = 0.406, *d* = 0.25.

**Figure 1.**
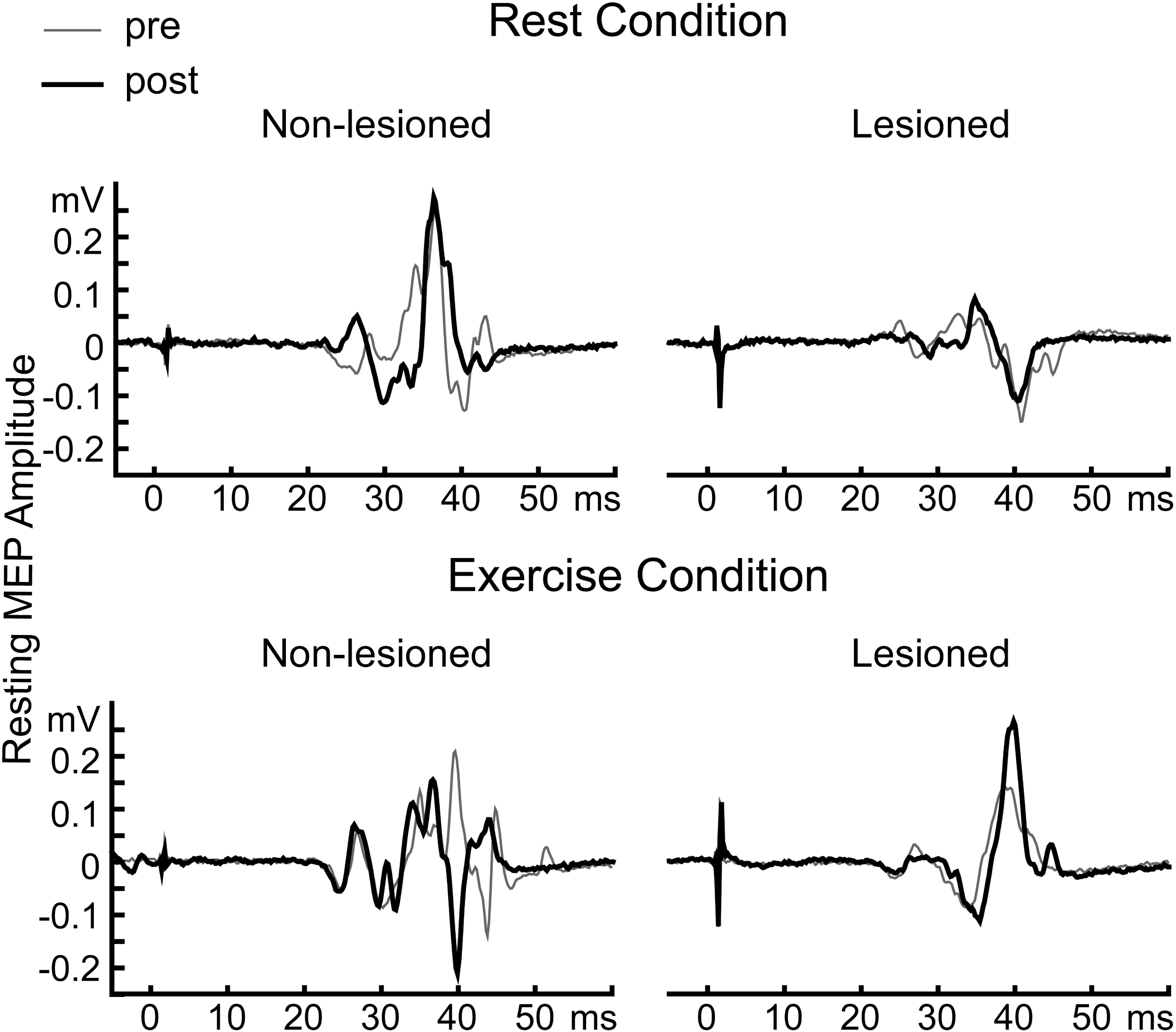
Raw EMG showing resting MEP traces (TMS intensity, 120% RMT) from an exemplar participant. Thin gray traces show pre-exercise or rest MEPs; thick black traces show post-exercise or rest MEPs. MEPs from the lesioned hemisphere increased after exercise, while they did not change after rest. MEPs from the non-lesioned hemisphere did not change after either exercise or rest. X-axis show time from TMS stimulus.

**Figure 2.**
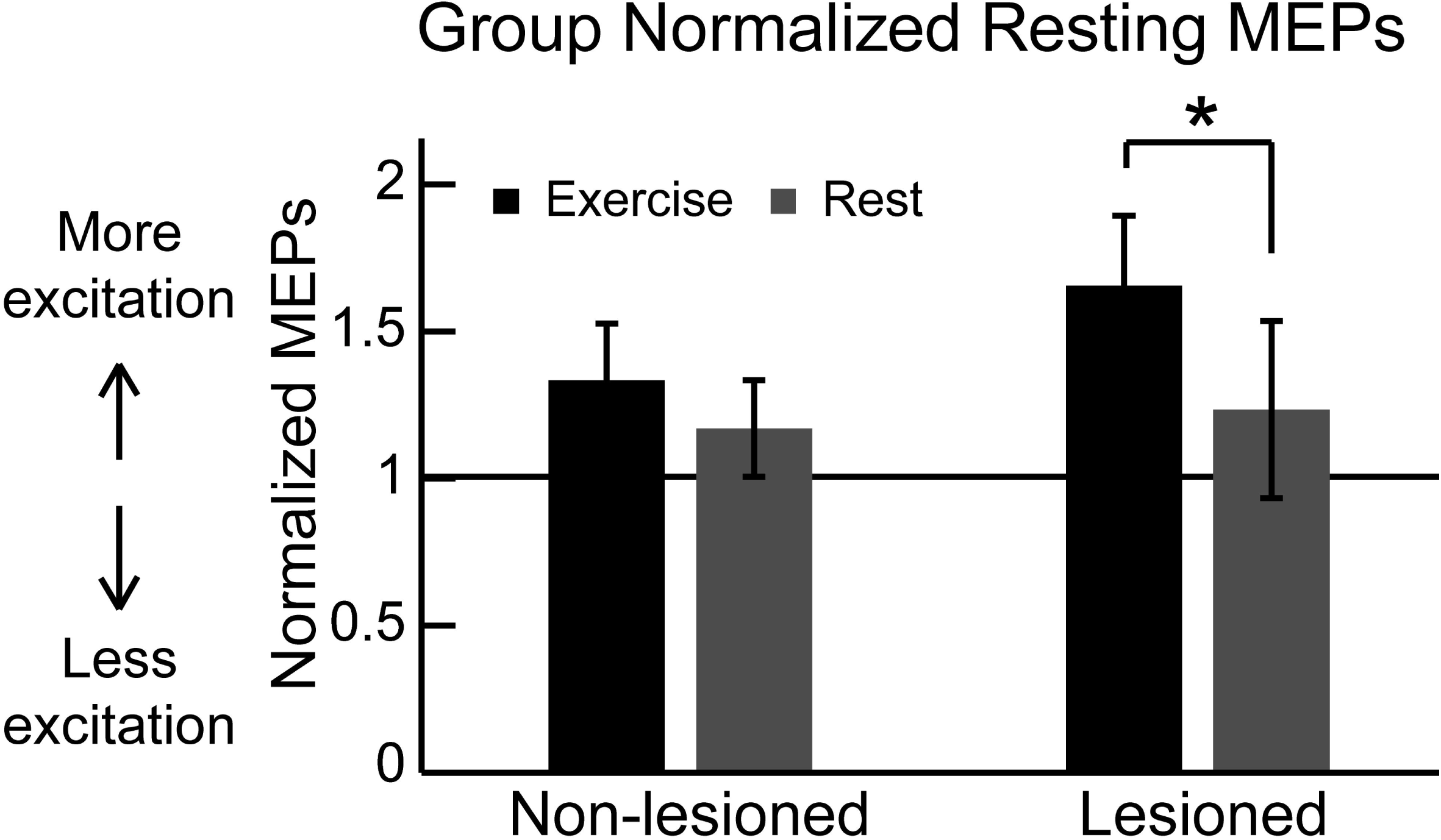
Group normalized (post/pre ratio) resting MEP amplitudes. Data from the exercise session are in black bars; data from the rest session are in gray bars. Value of 1 means no changes in MEP amplitude after exercise or rest. *Indicates significance between exercise vs. rest sessions at *p* < 0.05.

### SICI changes after exercise

SICI was also not significantly different before and after exercise in either the non-lesioned (pre, 0.47 ± 0.13; post, 0.55 ± 0.19) or the lesioned hemisphere (pre, 0.55 ± 0.19; post, 0.50 ± 0.12), both *p* > 0.60.

## Discussion

Here, we have shown that a short bout of high-intensity lower extremity exercise can increase motor cortical excitability in non-exercised muscles of people post-stroke. Specifically, peak-to-peak resting MEP amplitudes from the lesioned hemisphere were increased in this population following exercise priming. Notably, this is the first study to definitively detect positive effects of a clinically feasible (i.e., short) exercise priming protocol on motor cortical excitability in stroke.

Exercise has widespread positive influences on multiple levels of brain function^8^, some of which may explain the mechanisms underlying our results. Exercise can enhance long-term potentiation (LTP) in mice^32^, as well as up-regulate genes associated with the excitatory glutamatergic system, while down-regulating genes associated with the inhibitory GABA system in rats^33^. In healthy humans, acute exercise has been shown to enhance the effects of neuromodulation protocols that produce neuroplastic changes in the brain^34,35^. In addition, lactate uptake in the brain increases with high-intensity exercise^36,37^, and blood lactate increases, with or without exercise, have been associated with increased motor cortical excitability in healthy individuals^38^. This could explain why the high-intensity exercise used in our study may be effective in inducing motor cortical excitability changes. A neurochemical model has also been proposed to explain the effects of exercise on motor cortical excitability through modulation of arousal-related neurotransmitters, including dopamine, serotonin, and norepinephrine^39^. Brain-derived neurotrophic factor (BDNF) is also a potential candidate for increasing motor cortical excitability after exercise. Peripheral BDNF levels increase after exercise^40–43^ and are associated with LTP in both the hippocampus^44^ and the motor system^45^. However, interpretation of peripheral values of BDNF calls for caution^46^, and a recent study did not find increases of BDNF after exercise, nor was there a correlation between BDNF levels and acute exercise-induced neuroplasticity^35^. In summary, the mechanisms with which exercise induces neuroplasticity are unclear and need to be studied more systematically in the future.

Most exercise priming studies in healthy adults have reported increases in SICI following exercise, but have failed to detect changes in MEP amplitudes^9–11^. Several factors may have contributed to the difference between our results and these healthy cohort studies. Most obviously, the presence of a brain lesion in our participants with stroke would be expected to potentially influence brain responses to acute exercise. Interestingly, MEP amplitudes from the healthier non-lesioned hemisphere were not affected by exercise in our study, which is consistent with results from most healthy cohort studies. Second, aging can alter motor cortical excitability^47^, and change responses to other neuroplasticity-inducing protocols^48,49^. Thus it is possible that the older age of our participants may have affected the type and/or degree of plastic changes exhibited within the motor cortex in response to acute exercise. Last, differences in exercise paradigms may also have affected the results. We purposefully chose our exercise paradigm to be minimally-fatiguing so as to be easily administered clinically for people with stroke^20^. The fact that just 5 minutes of high-intensity exercise was able to induce resting MEP changes in the lesioned hemisphere is evidence for the potential of this paradigm to be used as an adjunct to physical therapy in the clinic, as a means to induce increased cortical excitability in patients with stroke.

In people with stroke, there are currently few other reports of excitability changes induced through exercise priming. More importantly, these studies all used longer, more fatiguing exercise protocols, which are not suitable for use in the clinic in conjunction with physical therapy. Using low-intensity exercise, Murdoch et al^50^ did not find any effect of exercise on motor cortical excitability post-stroke. Likely the exercise intensity used was insufficient to drive motor cortical excitability changes. In two other studies utilizing high-intensity exercise, one study found that a graded exercise test modulated the ratio of SICI levels between hemispheres but did not change SICI levels within a hemisphere nor did it change MEP amplitudes^18^. The other study utilized high-intensity interval training and found that MEP latencies from the lesioned hemisphere were lengthened after exercise, but intracortical measurements were unaffected^19^. MEP amplitudes were not reported. The exercise parameters used and the impairment levels of the participants may explain the differences in our results. Our exercise protocol was shorter and our stroke participants were on average more severely impaired. Perhaps, surprisingly, more severely impaired individuals may show an enhanced capacity to benefit from exercise priming. Regardless, the facts that our participants experienced a significant increase in lactate levels after exercise and had a positive excitability improvement in the lesioned hemisphere suggests that 5 minutes of fast walking can be sufficient to induce exercise priming effects in this population.

Interestingly, in the current study, MEP amplitudes in the non-lesioned hemisphere failed to show the same increases following exercise that were robust in the lesioned hemisphere. This result is somewhat surprising because one might presume that the lesioned hemisphere would be less responsive to changes induced by interventions, especially in the chronic stage of stroke. However, Abraha et al^19^ also found exercise-induced changes only in the lesioned hemisphere. Thus, it appears that post-stroke, the lesioned hemisphere may in fact be more predisposed to acute exercise priming than the non-lesioned hemisphere. This would be an exciting finding, because the ability to increase MEP amplitudes in the lesioned M1 is associated with better motor recovery^5–7^. Thus, acute exercise may be a potential method to promote better motor recovery post-stroke.

Our other measurement, SICI, was unaffected by acute exercise in either hemisphere. SICI is a measure of GABA_A_-mediated intracortical inhibition^51,52^. Based on our results, this type of intracortical inhibitory circuits may be less susceptible to changes induced by a short bout of exercise in people with stroke.

Some limitations to this work should be mentioned. First, because of concerns over session length and the probability of obtaining measures from the lesioned hemisphere, we opted not to measure intracortical facilitation or interhemispheric inhibition, both of which have been shown to change after acute exercise in healthy individuals^10,11^. Whether or not our priming paradigm affects these measures should be investigated. Also, it is important to note that we do not know whether the positive motor cortical excitability changes we found would be associated with gains in motor performance or learning. Nepveu et al^17^ did find better retention of a time-on-target upper extremity motor task after high-intensity interval training in people with stroke, but other work using a different learning paradigm suggests this effect may be very task-specific^53^. It remains to be investigated whether our exercise priming protocol can enhance motor performance or learning in people with stroke.

In conclusion, we found that just 5 minutes of high-intensity lower extremity exercise in people with stroke significantly increases motor cortical excitability within the lesioned hemisphere as measured by resting MEP amplitudes. Future studies should investigate whether this priming effect in the brain can be utilized in therapeutic interventions to improve motor recovery post-stroke.

## Geolocation Information

Newark, Delaware 19713

## Acknowledgments

Funding: This work was supported by the Delaware Economic Development Office under grant number 16A00377 and the National Institutes of Health under grant numbers S10RR028114, 1R01HD078330, and P20GM103446.

